# Mechanochemical coupling tunes robustness of PAR polarity across developmental contexts in the *C. elegans* embryo

**DOI:** 10.64898/2026.04.09.717607

**Authors:** Ryunosuke Saito, Sungrim Seirin-Lee, Masatoshi Nishikawa

## Abstract

Proper establishment of cell polarity is essential for reliable asymmetric divisions during embryonic development. In the *Caenorhabditis elegans* zygote, polarity is established through mechanochemical coupling between advective transport driven by cortical flows and mutually antagonistic PAR protein interactions. How these mechanisms are integrated across developmental contexts to establish cell polarity in the early embryo remains unexplored. Here we show that P_1_ polarization relies primarily on sustained antagonistic activity of anterior PAR proteins mediated by CDC-42 and PKC-3. Cortical contractility is regulated by CDC-42 through MRCK-1 to drive cortical flow, and is largely independent of the RHO-1 – LET-502 pathway that is crucial in zygotic polarity establishment. We find that these flows contribute only weakly to P_1_ polarity and primarily during the late phase. P_1_ polarization is sensitive to CDC-42 dosage, as indicated by reduced cytoplasmic asymmetry of MEX-5 and loss of division asynchrony between daughter cells. In contrast, zygotic polarization remains robust to comparable perturbations in CDC-42 expression when cortical flow is intact, but becomes sensitized when cortical flow is suppressed. Together, these findings show that cortical flow acts as a transport-mediated reinforcement that buffers PAR polarity against perturbations in CDC-42 expression, and that weakening this reinforcement sensitizes polarity establishment across developmental contexts.

## Introduction

The partitioning-defective (PAR) protein network plays an important role in regulating cell polarity across diverse biological contexts, and is evolutionarily conserved in *Caenorhabditis elegans, Drosophila melanogaster*, and many vertebrates[1, 2, 3]. The PAR network establishes mutually exclusive cortical domains that break the cellular symmetry and generate a polarized state. *C. elegans* zygote is a well-established model for studying PAR polarization. Two mutually exclusive domains, anterior and posterior PARs (aPAR and pPAR), define the anterior–posterior axis of the embryo, and the boundary between them determines the position of the future division plane, producing daughter cells with distinct sizes and developmental fates[4, 5, 6, 7, 8].

Initially, the cortex of the zygote is occupied by the aPAR complex, composed of the atypical protein kinase PKC-3, the adapter PAR-6, and the cortical scaffolds PAR-3 or CDC-42[9, 10, 11, 12]. The sperm-donated centrosome, typically positioned at the future posterior, locally reduces non-muscle myosin II (NMY-2) driven contractility, thereby establishing spatial gradients of cortical tension that drive a large-scale anterior-directed cortical flow[13, 14, 15]. This flow transports the PKC-3–PAR-6–PAR-3 complex toward the anterior, initiating segregation of aPAR proteins[14, 16, 17, 18, 19]. The centrosome also plays an initiating role by preventing PAR-2 from being phosphorylated by PKC-3, thereby promoting its loading onto the posterior cortex and facilitating the initiation of pPAR domain formation, even in the absence of cortical flow[20, 21]. After polarity is established, it is maintained by mutually inhibitory interactions between aPAR and pPAR proteins. CDC-42 plays an indispensable role in this process, as its depletion disrupts aPAR segregation and causes spreading toward the posterior[12, 22, 23, 24, 25, 26]. PKC-3 bound to CDC-42 phosphorylates the posterior PAR proteins PAR-1 and PAR-2, promoting their dissociation from the cortex[27, 28, 29, 30, 31], while PAR-1 phosphorylates PAR-3 to exclude it from the cortex[21]. Notably, although neither advective transport nor mutual inhibition triggered by PAR-2 loading is strictly essential, either mechanism alone can establish polarity, underscoring the robustness of zygotic polarization to perturbations in protein concentrations[5, 16, 17, 32].

P_1_ cell, a daughter cell of zygote, re-establishes polarity to undergo a subsequent asymmetric division. Previous studies have shown that P_1_ polarization relies on the interplay between advective transport and biochemical reactions, as in the zygote[33, 34]. However, cortical flow in P_1_ polarization arises only in the later phase and at a reduced rate[14, 33, 34, 35], in sharp contrast to the zygotic polarization, which is initiated by prominent long-ranged flows. This raises the question of how advective transport and biochemical reactions are differently coupled during P_1_ cell polarization compared to the zygote. It therefore remains unclear how robust polarity establishment remains when mechanochemical coupling between cortical flow and biochemical reactions is reduced.

Here we show that antagonistic activity by anterior PARs, mediated by PKC-3 and CDC-42, is required throughout P_1_ polarity establishment, whereas advective transport driven by cortical flow contributes only during the late phase and plays a minor role. We further show that CDC-42 regulates cortical contractility through MRCK-1 to generate cortical flows that advect aPARs. P_1_ polarization is sensitive to changes in CDC-42 expression level, whereas P_0_ polarization remains robust when advective transport driven by cortical flow is intact. Our results reveal that mechanochemical coupling between advective transport and antagonistic interactions tunes the robustness of PAR polarity across developmental contexts.

## Results

### Characterizing PAR polarization dynamics

To investigate the mechanochemical coupling during P_1_ polarization, we first sought to characterize the PAR polarization dynamics. Immediately following completion of cytokinesis, PAR-2 intensity at the anterior cortex of the P_1_ cell exhibited a rapid and substantial decrease within approximately 4 min (Fig. 1A and C). This decrease occurred during an early phase of polarization, in contrast to the about 11 min cell-cycle duration of the AB cell [34]. Based on these data, we conclude that PAR-2 polarity is largely established during the early phase of P_1_ polarization.

**Figure 1.**
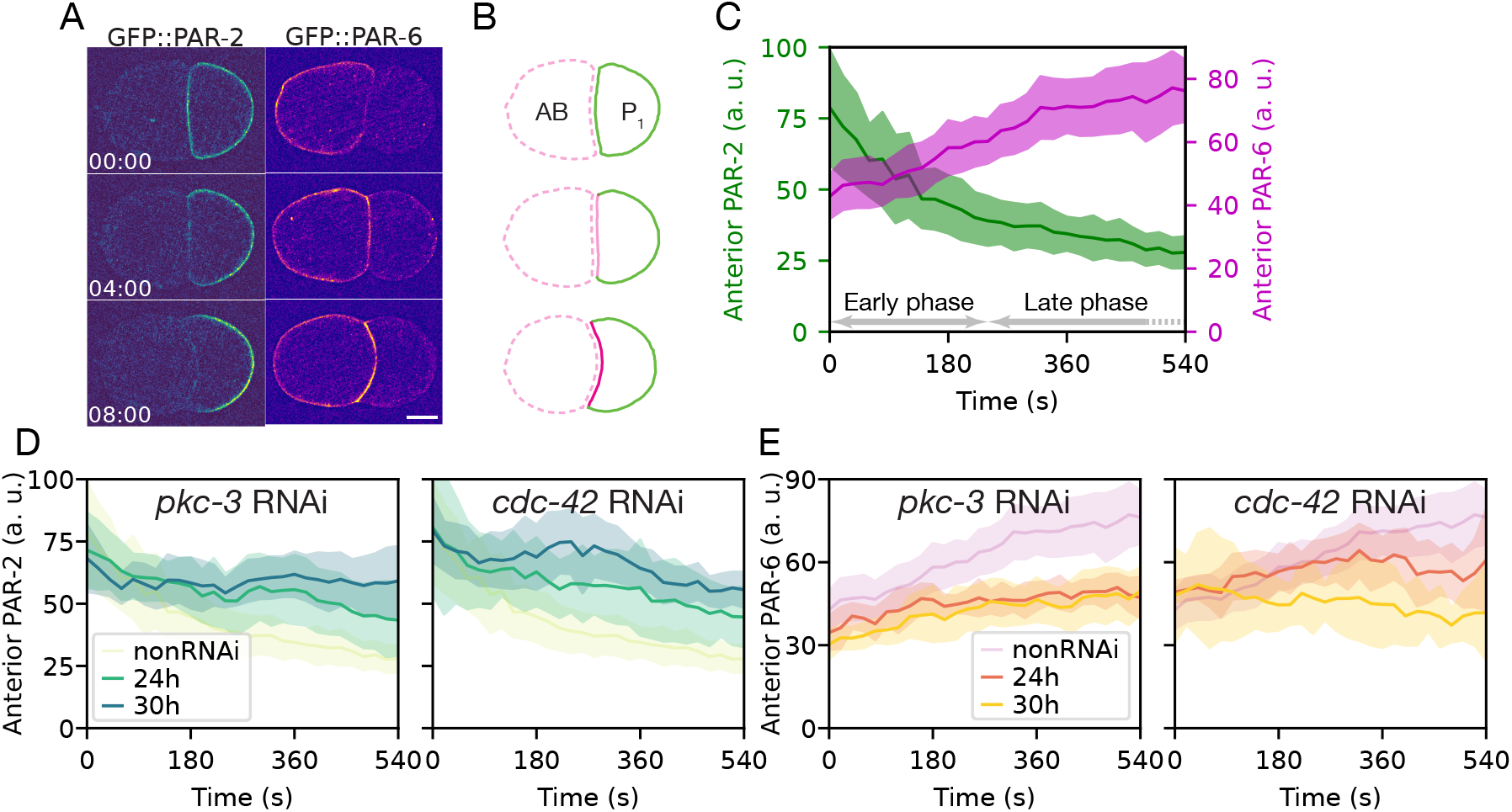
Anterior PAR antagonism is required throughout P_1_ cell polarization. (A) Representative images of posterior PAR (pPAR, images of PAR-2::GFP in left) and anterior PAR (aPAR, images of PAR-6::GFP in right) during P_1_ cell polarization. Scale bar, 10*µ*m. (B) Schematic of P_1_ cell polarization, in which pPAR is cleared from the anterior cortex while aPAR accumulates there over time. (C) Time-courses of fluorescence intensities of pPAR (n = 9) and aPAR (n = 14) at the anterior cortex of P_1_ cell. Shaded regions denote 95 % confidence intervals of the mean. The transition from an early to a late polarization phase is indicated. (D-E) Mild RNAi perturbations targeting *pkc-3* (left) and *cdc-42* (right), slow P_1_ cell polarization dynamics. Partial depletion of either factor delays posterior PAR clearance (D, *pkc-3* RNAi: n = 11 for 24 h, n = 10 for 30 h, *cdc-42* RNAi: n = 7 for 24 h, n = 9 for 30 h) and anterior PAR accumulation (E, *pkc-3* RNAi: n = 8 for 24 h, n = 9 for 30 h, *cdc-42* RNAi: n = 7 for 24 h, n = 6 for 30 at the anterior cortex across the polarization period.

We next examined aPAR polarization dynamics at the anterior cortex of the P_1_ cell. PAR-6 intensity at the AB/P_1_ contact increased gradually for approximately 9 min following completion of P_0_ cytokinesis, in contrast to the rapid decrease of PAR-2 during the first 4 min (Fig. 1A and C). Because optical microscopy cannot resolve two opposing cortical domains at cell-cell contacts[9, 33, 34], we tested whether this increase could arise from redistribution of PAR-6 within the AB cell. Neither cytoplasmic PAR-6 levels in AB nor PAR-6 intensity at the anterior AB cortex away from the AB/P_1_ contact site changed during polarization (Fig. S1A and B), demonstrating that posterior cortical PAR-6 concentration in AB cell remained unchanged. In contrast, the P_1_ cell exhibited a decrease in cytoplasmic PAR-6 and a transient increase in posterior cortical PAR-6[34] (Fig. S1A and B). Together, these observations support the conclusion that the increase in PAR-6 signal at the AB/P_1_ contact reflects accumulation at the anterior cortex of the P_1_ cell.

### aPAR antagonistic activity is essential for entire period of polarization

We next focus on understanding the mechanisms that establish P_1_ polarity by modulating antagonistic activity of anterior PAR proteins. To perturb P_1_ polarization dynamics while maintaining zygotic polarity, we applied mild genetic perturbations that minimally affect P_0_ polarization but influence subsequent polarization dynamics. Previous studies have pursued this goal using temperature-sensitive and analog-sensitive mutants [33, 34].

Here, we instead employed RNA interference targeting short gene fragments to achieve reproducible partial depletion. Using this approach, we generated feeding RNAi clones targeting *pkc-3* and *cdc-42* (Fig. S2A). These RNAi treatments had only minor effects on zygotic polarization and asymmetric division (Fig. S2B-D), enabling characterization of their contribution to P_1_ cell polarization dynamics.

Previous studies demonstrated an indispensable role for PKC-3 inhibitory interactions with posterior PAR proteins during P_1_ polarity establishment [33, 34]. We therefore investigated how partial depletion of PKC-3 affects P_1_ polarization dynamics. PKC-3 depletion slowed PAR-2 polarization throughout the entire period of P_1_ polarization (Fig. 1D left). Similarly, the rate of PAR-6 enrichment at the anterior cortex was reduced in partially PKC-3 depleted embryos (Fig. 1E left). Notably, PKC-3 depletion also attenuated the transient accumulation of PAR-6 at the posterior cortex during the early phase of polarization (Fig. S2G), indicating that early aPAR cortical recruitment at posterior also depends on PKC-3 activity. Taken together, we conclude that PKC-3 plays an indispensable role throughout the entire period of P_1_ cell polarity establishment.

We next asked whether CDC-42 is required for P_1_ polarity establishment. In the zygote, PKC-3 kinase activity that promotes dissociation of posterior PAR proteins requires complex formation with CDC-42 and PAR-6, and depletion of CDC-42 leads to diffusive spreading of anterior PAR proteins from the anterior domain[12]. Consistent with this role, partial depletion of CDC-42 in P_1_ slowed both PAR-2 clearance and PAR-6 accumulation at the anterior cortex throughout the entire period of P_1_ polarization (Fig. 1D right and E right), similar to the effects observed under partial *pkc-3* RNAi. Taken together, these data indicate that aPAR antagonistic activity mediated by the PKC-3–CDC-42 complex is indispensable throughout P_1_ polarity establishment.

### Advection plays only a limited role in P_1_ polarity establishment

We next examined cortical myosin dynamics during P_1_ polarization. During the late phase of polarization, cortical myosin flow in the P_1_ cell exhibited a significant posterior-to-anterior component with a peak velocity of −1.34 ± 0.449 *µ*m min^−1^(Fig. 2B and C), consistent with previous reports [14, 33, 35]. This flow coincided with a marked increase in cortical myosin density, most prominently in the AB cell, suggesting synchronous activation of cortical contractility in both daughter cells (Fig. S3 B). This late-phase increase in cortical myosin density was accompanied by cell rounding, a proxy for increased cortical contractility. Sectional curvature at the AB/P_1_ contact increased beginning 4 min after cytokinesis completion (Fig. S3 B). Together, these results indicate that late-phase cortical contractility is synchronously up-regulated in AB and P_1_ cells, driving cortical flow in P_1_ and rounding of the AB cell.

**Figure 2.**
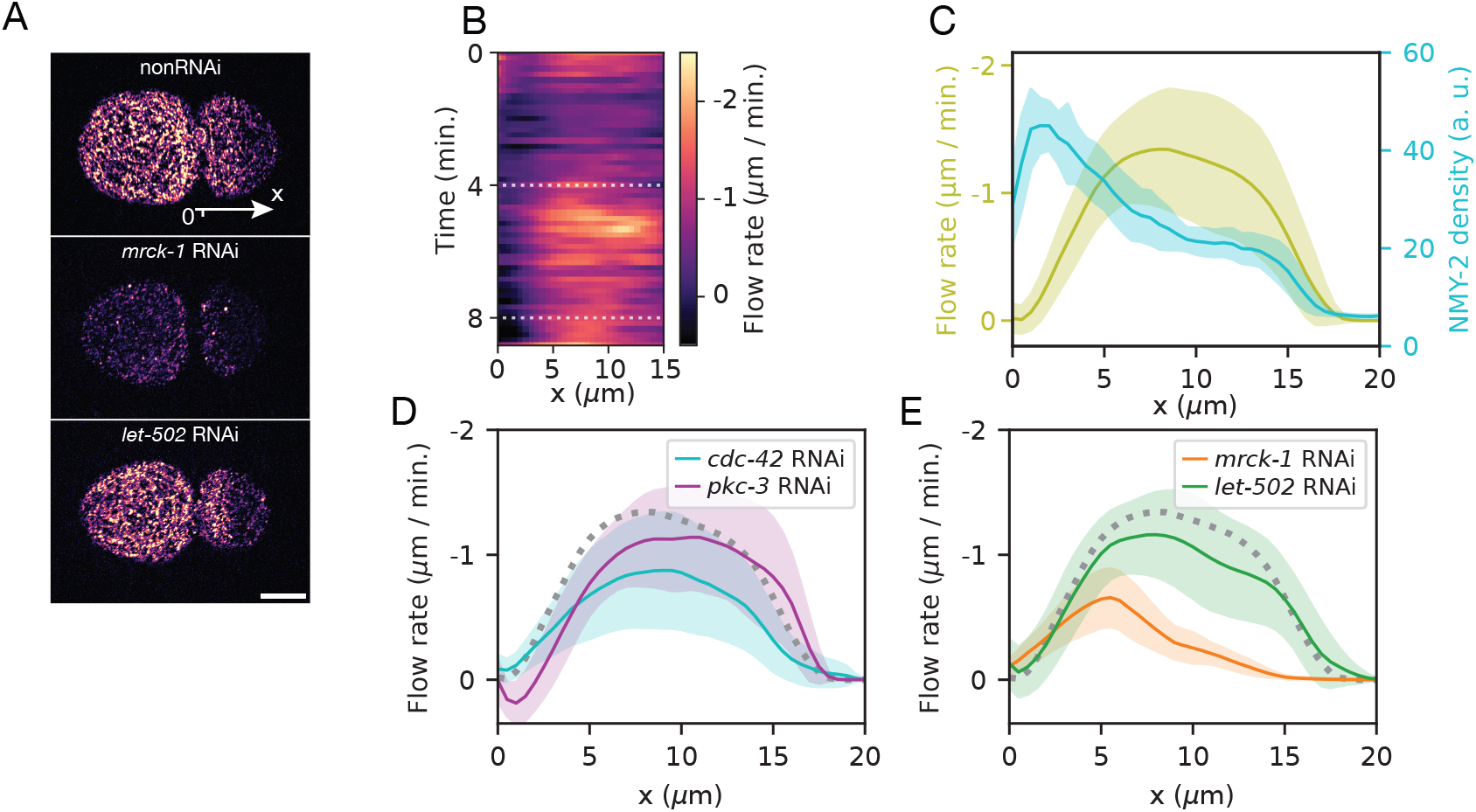
CDC-42 regulates cortical contractility and flow via MRCK-1. (A) Representative cortical NMY-2::GFP images at 2-cell stage from nonRNAi embryos and embryos subjected to RNAi against *mrck-1* and *let-502*. Scale bar, 10*µ*m. (B) Spatiotemporal profile of cortical flow rate along anterior-posterior (AP) axis of the P_1_ cell in nonRNAi embryos, (n = 11). Time T = 0 corresponds to completion of P_0_ cell cytokinesis. (C) Spatial profiles of NMY-2::GFP intensity and cortical flow rate along the AP axis of P_1_ cell during the late phase of polarization, ensemble averages were taken for temporal average profile between T = 4 min and T = 8 min (n = 11). Shaded regions indicate 95 % confidence intervals of the mean. (D-E) Spatial profiles of NMY-2::GFP intensity and cortical flow rate under RNAi perturbations targeting *cdc-42* and *pkc-3* (D, n = 10 and 10, respectively), *mrck-1* and *let-502* (E, n = 10 and 12, respectively).

While imaging PAR polarization, we noticed that the rounding up of AB cell in the late phase was attenuated in *cdc-42* but not in *pkc-3* RNAi embryos (Fig. S2 E, F and Fig.S3C), suggesting that CDC-42 regulates late-phase cortical contractility. Consistently, partial depletion of CDC-42 abolished the late-phase increase in cortical myosin density in both the AB and P_1_ cells (Fig. S3B). As a consequence, anterior-directed cortical flow in the P_1_ cell was significantly reduced, reaching a peak velocity of −0.875 ± 0.462*µ*m min^−1^, approximately 65 % of nonRNAi condition (Fig. 2D).

Given that CDC-42 regulates cortical contractility through MRCK-1 for zygotic polarity maintenance and gastrulation[26, 36], we asked whether CDC-42 mediated cortical regulation involves MRCK-1. *mrck-1* RNAi abolished the late phase increase of cortical NMY-2 density in both AB and P_1_ cells. Accordingly, peak cortical flow observed in late phase P_1_ cell was decreased to −0.657 ± 0.232 *µ*m min^−1^(Fig. 2E). Because MRCK-1 is expected to act downstream of CDC-42, we used a stronger *mrck-1* RNAi treatment than those applied to *cdc-42* or *pkc-3*. Notably, this treatment had minor effects on zygotic polarization, allowing us to test whether MRCK-1 is required for the generation of cortical flow during P_1_ polarization (Fig. S2D). Taken together, these results suggest that CDC-42 regulates cortical contractility and associated cortical flows during P_1_ polarity establishment through MRCK-1.

We next asked whether RHO-1 contributes to contractility regulation during P_1_ polarization. In the zygote, RHO-1 is a key regulator of cortical contractility that drives cortical flow during polarity establishment. To test whether this pathway also operates in the P_1_ cell, we examined cortical flow in embryos depleted of LET-502, the Rho-associated kinase phosphorylating the regulatory light chain of non-muscle myosin[37]. Under *let-502* RNAi, cortical flow in the P_1_ cell remained largely preserved, with a peak velocity of −1.16±0.338 *µ*m min^−1^during the late phase (Fig. 2E), in marked contrast to the sharp reduction observed in *mrck-1* RNAi embryos. These observations suggest that late-phase cortical contractility during P_1_ polarization is regulated primarily through the CDC-42–MRCK-1 pathway rather than through the canonical RHO-1–LET-502.

We then asked whether PAR polarity is required for cortical flow during the late phase of P_1_ polarization. In the zygote, *par* mutants, in which pPAR is homogeneously distributed on the cortex, fail to generate long-ranged flow[38, 14]. To test if the cortical flow in P_1_ cell is similarly influenced by PAR polarity, we investigated cortical flow in *pkc-3* RNAi embryos. Peak cortical flow observed in late phase P_1_ cell was −1.14 ± 0.416 *µ*m min^−1^(Fig. 2D), in sharp contrast to the reduction observed in *cdc-42* partial RNAi treated embryos and in the zygote under PAR polarity disruption. These results suggest that late-phase cortical flow in the P_1_ cell does not require proper PAR polarity establishment.

To dissect the mechanochemical interplay for polarity establishment, we then characterized advective transport of PAR protein complex driven by cortical flow. We tracked individual PAR-3::GFP clusters in the cortical plane using time-lapse imaging. In non-RNAi embryos, biased movement toward anterior was evident from the displacement trajectories and their temporal average (Fig. 3B). The average velocity along AP axis was −0.810 ± 0.211*µ*m min^−1^, comparable to the cortical flow rate determined above (Fig. 3C). In contrast, strong depletion of MRCK-1 reduced the mean AP velocity to be −0.198 ± 0.159 *µ*m min^−1^(Fig. 3B right and C). This reduction was consistent with the corresponding attenuation of cortical flow. Together, these results indicate that anterior-directed transport of PAR-3 during the late phase of P_1_ polarization is driven primarily by cortical flow regulated through MRCK-1. Consistent with this, reduced PAR-3 advection was also observed in *cdc-42* RNAi embryos, whereas *pkc-3* RNAi embryos retained anterior-biased motion (Fig. S5).

**Figure 3.**
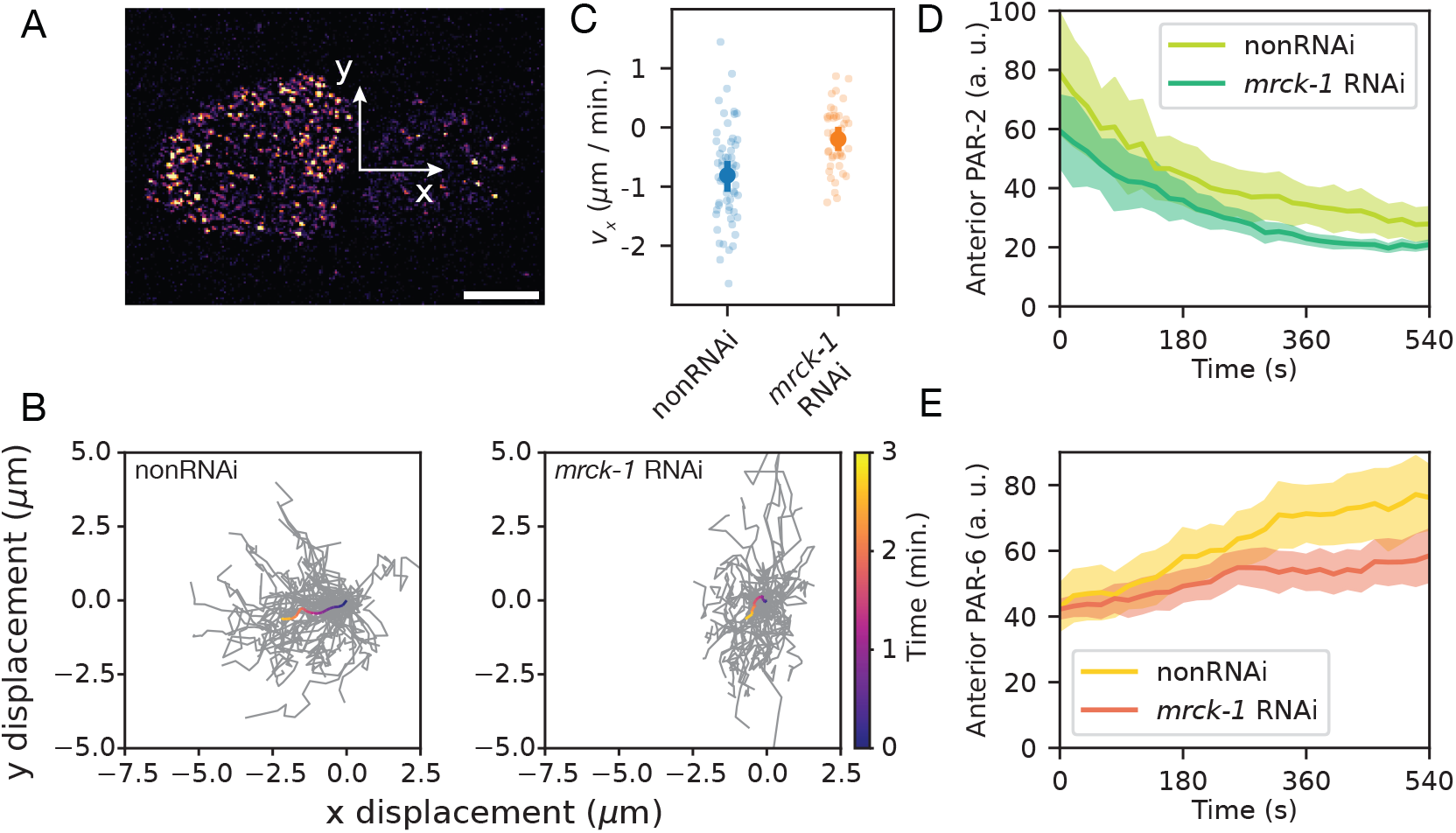
Cortical flow advects PAR-3 clusters via CDC-42 and MRCK-1 during P_1_ polarization. (A) Representative cortical PAR-3::GFP images at 2-cell stage. Scale bar, 10*µ*m. (B) Trajectories of individual PAR-3 clusters during the polarization period. Grey lines show trajectories of individual clusters, and temporally color-coded lines indicate ensemble-averaged trajectories over time. Data are shown for nonRNAi embryos (left, n = 57 clusters from 8 embryos) and *mrck-1* RNAi (right, n = 45 clusters from 9 embryos). (C) Distributions of PAR-3 cluster velocities along the AP axis for the indicated conditions, computed from individual cluster trajectories shown in (B). (D-E) Time courses of pPAR (D, n = 10), and aPAR (E, n = 16) fluorescence intensities measured at the anterior cortex of P_1_ cell. Shaded regions indicate 95 % confidence intervals of the mean.

To characterize the contribution of advective transport to P_1_ polarity establishment, we examined PAR polarization dynamics in *mrck-1* RNAi, which suppresses cortical flow. PAR-2 polarization dynamics was indistinguishable from those in nonRNAi embryos, in which the rapid and substantial decrease of PAR-2 just after the completion of preceding cytokinesis, prior to cortical flow[34] (Fig. 3D). Similarly, early-phase accumulation of PAR-6 at the anterior cortex was largely unchanged (Fig. 3E).

In contrast, PAR-6 polarization during the late phase was affected by *mrck-1* RNAi. The late-phase increase in PAR-6 intensity at the anterior cortex was abolished by MRCK-1 depletion (Fig. 3E), whereas nonRNAi embryos exhibited continued PAR-6 accumulation. These results demonstrate that advective transport contributes to PAR-6 accumulation at late phase of P_1_ polarization[33]. However, the overall contribution of advective transport to P_1_ polarity establishment is limited, as the early phase of polarization largely relies on antagonistic interactions.

### Polarization robustness

Given that advective transport plays a limited role in P_1_ polarity establishment, we asked whether P_1_ polarization is less robust to perturbations in polarity protein expression levels than zygotic polarization, in which redundancy between advective transport and antagonistic interactions ensures robust polarity establishment[17, 32]. To address this, we characterized cytoplasmic MEX-5 asymmetry between the two daughter cells (Fig. 4A), as well as the timing difference in nuclear envelope breakdown (NEBD) (Fig. 4D), as functional readouts of polarity-dependent cell-cycle asynchrony in embryos subjected to stronger CDC-42 depletion.

**Figure 4.**
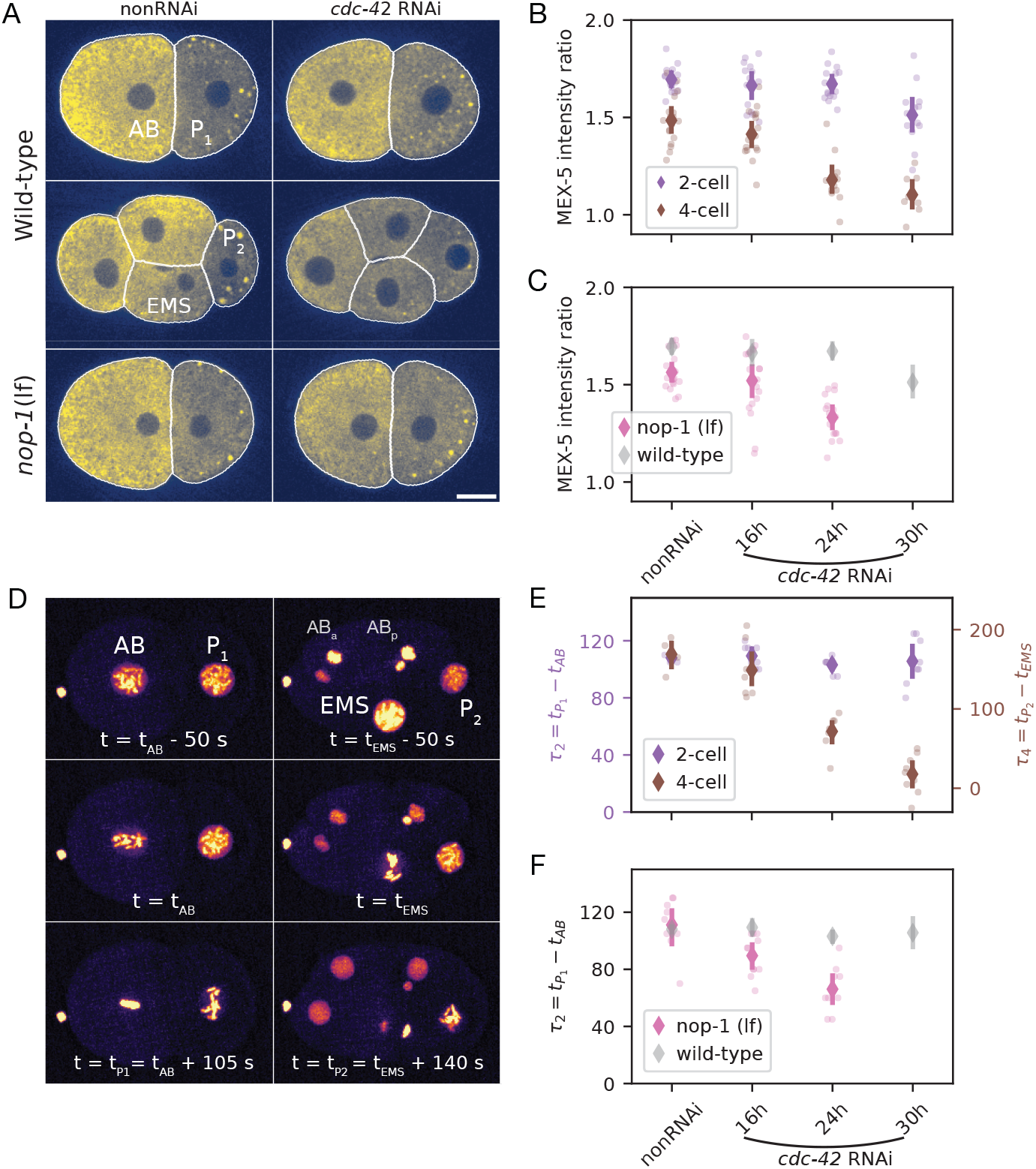
Sensitivity of cell asymmetry to CDC-42 expression level in wild-type and flow-deficient embryos. (A) Representative images of MEX-5::mNeonGreen (mNG) distributions in wild-type embryos at the 2-cell and 4-cell stages under nonRNAi and *cdc-42* RNAi conditions. White outlines indicate cell contours used for fluorescence quantification. Scale bar, 10*µ*m. (B) MEX-5 intensity ratios between sister cells at the 2-cell stage (AB/P_1_) and 4-cell stage (EMS/P_2_) in wild-type embryos subjected to graded *cdc-42* RNAi. Error bars represent 95 % confidence intervals of the mean. Ratios are averaged across n = 11 and 13 embryos (nonRNAi), n = 11 and 9 embryos (16 h RNAi), n = 10 and 10 embryos (24 h RNAi), n = 10 and 10 embryos (30 h RNAi), for 2-cell and 4-cell stages, respectively. (C) MEX-5 intensity ratios between AB and P_1_ cells in *nop-1 (lf)* mutant embryos, in which cortical flow is strongly suppressed. Wild-type data are shown in gray for comparison. Intensity ratios are averaged across 19 embryos (nonRNAi), 22 embryos (16 h RNAi), 20 embryos (24 h RNAi). (D) Representative timelapse images showing nuclear envelope breakdowns (NEBD) at 2-cell (left) and 4-cell stage (right). Time differences of NEBD between AB and P_1_ cells at the 2-cell stage, and EMS and P_2_ cells at the 4-cell stage, in wild-type embryos subjected to graded *cdc-42* RNAi. Error bars represent 95 % confidence intervals of mean. Data are averaged across n = 5 and 5 embryos (nonRNAi), n = 8 and 9 embryos (16 h RNAi), n = 8 and 9 embryos (24 h RNAi), n = 8 and 9 embryos (30 h RNAi), for 2-cell and 4-cell stages, respectively. Time differences of NEBD between AB and P_1_ cells in *nop-1 (lf)* mutant embryos under *cdc-42* RNAi. Data are averaged across 9 embryos (nonRNAi), 11 embryos (16 h RNAi), 9 embryos (24 h RNAi).

To investigate the cytoplasmic asymmetry, we determined the ratio of MEX-5 intensities between two sister cells produced by asymmetric cell divisions, for various RNAi feeding times. The MEX-5 intensity ratio between AB and P_1_ cells remained largely unchanged across RNAi feeding time ranging from 16 to 30 hours, and was comparable to that of nonRNAi embryos (Fig. 4B purple). Remarkably, the MEX-5 intensity ratio between P_2_ and EMS cells gradually decreased to approach unity with longer *cdc-42* RNAi treatment (Fig. 4B brown), suggesting a loss of cytoplasmic asymmetry in P_1_ cell. Notably, previous studies reported that stronger perturbations, such as longer RNAi treatment through injection, were required to induce symmetric division of P_0_ cell[22, 23].

MEX-5/6 are involved in asymmetric inheritance of polo-like kinase PLK-1 and cyclin-dependent kinase phosphatase CDC-25.1, which promote mitotic entry, and thus is expected to controls cell division asynchrony in two sister cells produced by asymmetric cell division[39, 40, 41]. We thus examined cell division asynchrony under *cdc-42* RNAi. To this end, we measured the time difference in nuclear envelope breakdown (NEBD) between sister cells following asymmetric divisions (Fig. 4D). At two cell stage, the NEBD timing difference, defined as 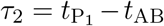, where 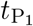 and *t*_AB_ denote the the times of NEBD in P_1_ and AB cells respectively, was not significantly altered compared to nonRNAi embryos (Fig. 4E purple). This indicates that division asynchrony at the two-cell stage is robust to the reduction in CDC-42 levels tested here, consistent with the preserved asymmetry of MEX-5 inheritance.

In contrast, division asynchrony at the four-cell stage was markedly affected. The NEBD timing difference, 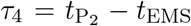, progressively decreased with longer *ccdc-42* RNAi treatment (Fig. 4E brown). After 30 h RNAi, a subset of embryos exhibited near-synchronous divisions of the P_2_ and EMS cells. Taken together, these results demonstrate that cell-division asynchrony arising from asymmetric inheritance of the cytoplasmic polarity mediator MEX-5 is less robust at the four-cell stage than at the two-cell stage with respect to perturbations in CDC-42 expression.

Finally, we next tested whether loss of cortical flow renders zygotic polarity establishment as sensitive to CDC-42 perturbation as that of the P_1_ cell. To address this, we characterized cytoplasmic asymmetry of MEX-5 under conditions in which cortical flow is strongly suppressed. *nop-1* is required for RHO-1 activation during polarity establishment to drive cortical flow, while contributing redundantly during cytokinesis [42, 43]. We therefore measured the cytoplasmic MEX-5 intensity ratio in *nop-1(it142)* mutant embryos subjected to strong *cdc-42* RNAi.

Under these conditions, the MEX-5 intensity ratio between the AB and P_1_ cells progressively decreased with longer RNAi treatment, in a similar manner to the ratio between EMS and P_2_ cells (Fig. 4 B, C). Consistent with this, the NEBD timing difference (*τ*_2_) also decreased with increasing RNAi duration (Fig. 4F). Both the MEX-5 intensity ratio and NEBD timing difference were largely unchanged in nonRNAi *nop-1 (it142)* embryos, indicating that the observed reduction of cytoplasmic asymmetry and division asynchrony reflects increased sensitivity to CDC-42 perturbation when cortical flow is suppressed. Together, these results demonstrate that suppressing cortical flow renders zygotic polarity establishment sensitive to perturbations in CDC-42 expression, recapitulating the reduced robustness observed in P_1_ polarization.

## Discussion

Our results reveal how polarity robustness arises from the mechanochemical coupling between biochemical antagonism and advective transport driven by cortical flow in the early C. elegans embryo. P_1_ polarization relies primarily on sustained antagonistic activity of anterior PAR proteins mediated by CDC-42 and PKC-3, whereas cortical flow arising in the late phase contribute only weakly. Reducing CDC-42 levels compromises MEX-5 asymmetry and division asynchrony in P_1_ descendants. Importantly, MEX-5 asymmetry and division asynchrony in P_0_ descendants remain robust to perturbations in CDC-42 expression levels only when advective transport driven by cortical flow is present. Together, these findings suggest that the extent of mechanochemical coupling tunes the robustness of PAR-driven polarity across developmental contexts.

We identify CDC-42 as a central regulator of the mechanochemical interplay during P_1_ polarization, acting not only through its canonical role in activating PKC-3 mediated antagonistic interactions but also by controlling cortical contractility via its effector kinase MRCK-1. In the zygote, RHO-1 primarily regulates cortical contractility that generates prominent long-range cortical flows facilitating polarization, whereas CDC-42 functions mainly to spatially restrict PKC-3 kinase activity during the maintenance phase. In contrast, during P_1_ polarization, cortical flows arise only later and at reduced strength, consistent with reduced inheritance of cortical myosin. As a result, advective transport provides only limited reinforcement of polarity establishment in P_1_, rendering the system more sensitive to perturbations of biochemical interactions, such as reductions in the expression levels of PAR proteins. Consistent with previous studies showing that cortical flow buffers PAR polarity in the zygote, our results extend this framework by demonstrating that loss or weakening of flow-mediated reinforcement renders polarity establishment sensitive to reductions in CDC-42 expression across developmental contexts. Altogether, our work reveals that developmental robustness of PAR polarity emerges from mechanochemical coupling and can be lost when this coupling is weakened or altered across developmental contexts.

---

**Figure S1:**
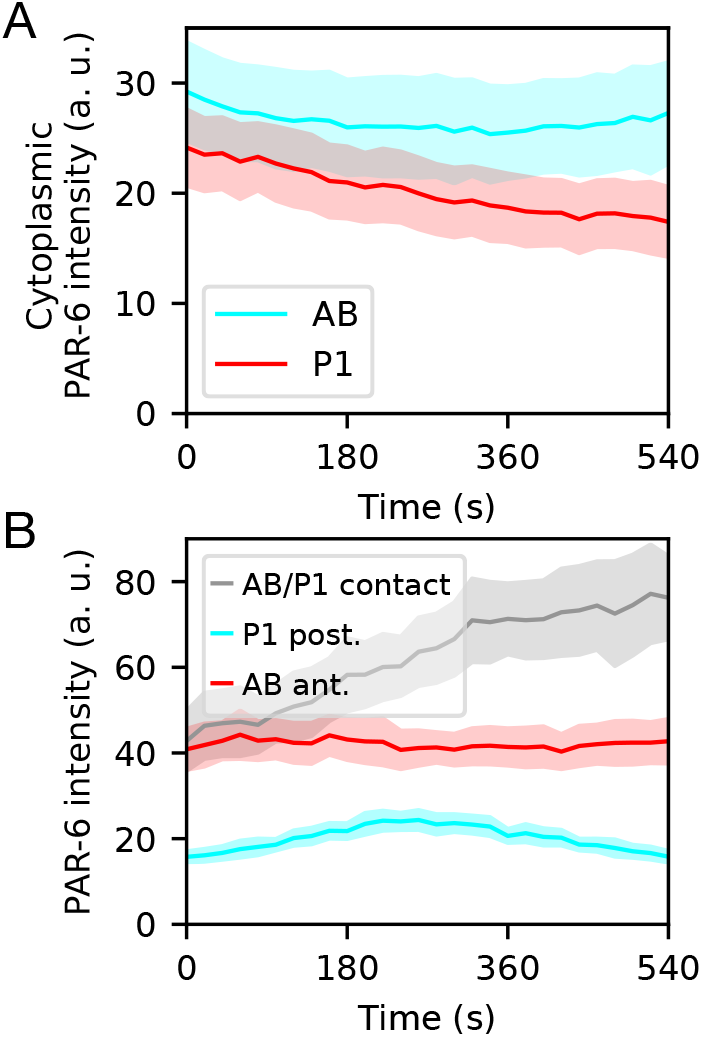
Cortical PAR-6 distribution in AB/P_1_ cells during P_1_ polarization. (A) Cytoplasmic PAR-6 intensities in the AB (cyan) and P_1_ (red) cells. (B) Cortical PAR-6 distribution at the anterior of AB and posterior of P_1_, away from the cell contact in which PAR-6 accumulation occurred, in the mid-plane of the embryo.

**Figure S2:**
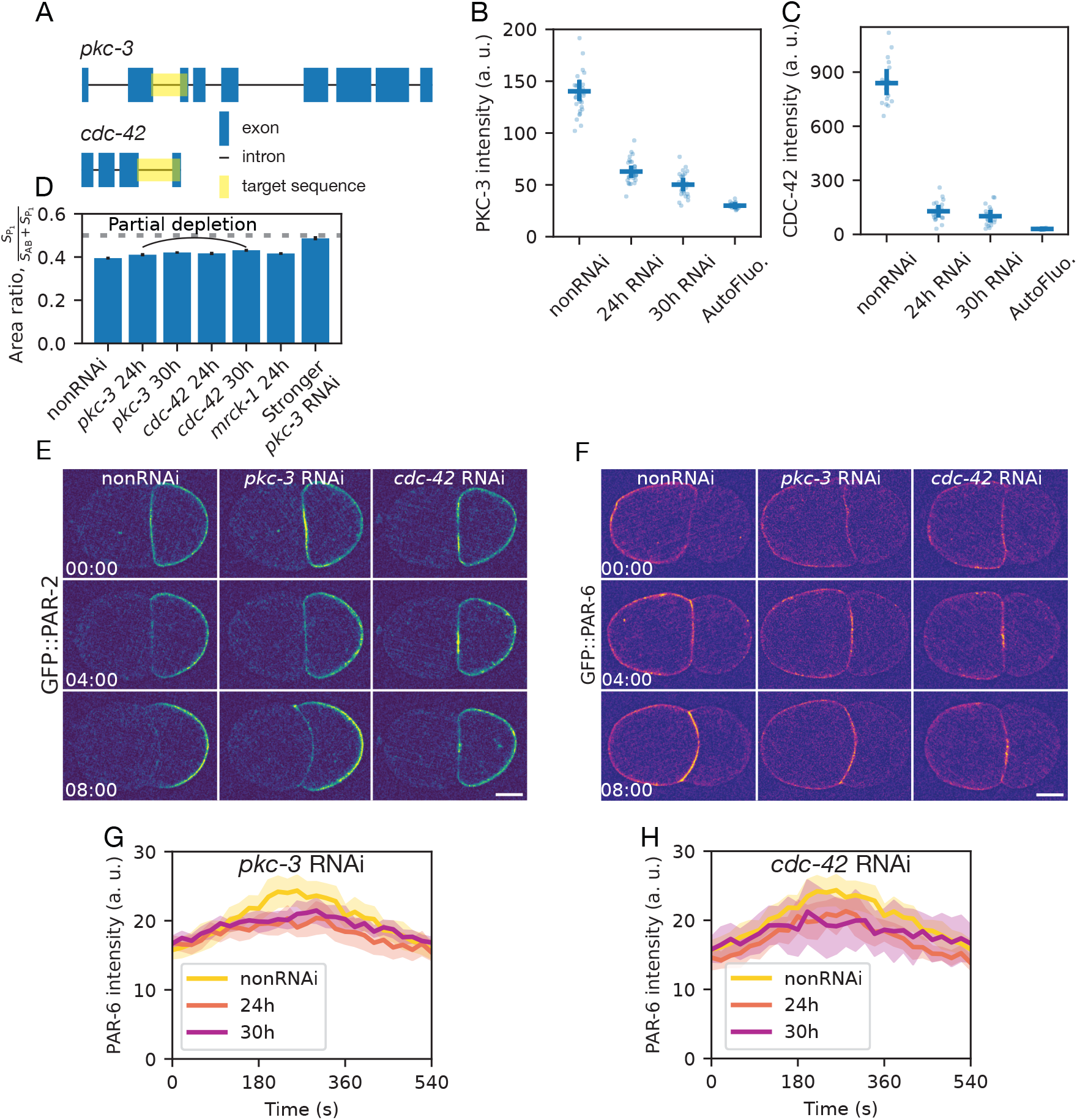
Mild RNAi perturbations targeting *pkc-3* and *cdc-42*. (A) Schematic of the target sequences for *pkc-3* and *cdc-42*, respectively. (B-C) Detected intensities of the GFP fused PKC-3 (B) and CDC-42 (C) subjected to mild RNAi targeting *pkc-3* and *cdc-42*, respectively. Error bars indicate 95 % confidence intervals of the mean. In (B), n = 30 (nonRNAi), n = 30 (24 h RNAi), n = 26 (30 h RNAi). In (C), n = 17 (nonRNAi), n = 21 (24 h RNAi), n = 20 (30 h RNAi). (D) Cell area ratio measured from mid-plane images. n = 10 (nonRNAi), n = 11 (*pkc-3* RNAi 24 h), n = 10 (*pkc-3* RNAi 30 h), n = 11 (*cdc-42* RNAi 24 h), n = 12 (*cdc-42* RNAi 30 h), n = 10 (*mrck-1* RNAi 24 h), n = 10 (stronger *pkc-3* RNAi). Error bars indicate 95 % confidence intervals of the mean. (E-F) Polarization dynamics of PAR-2 (E) and PAR-6 (F) under mild RNAi targeting *pkc-3* and *cdc-42*. T = 00:00 corresponds to completion of P_0_ cytokinesis. Scale bar, 10*µ*m. (G-H) Time courses of posterior PAR-6 accumulation under mild RNAi treatments, targeting *pkc-3* (G) and *cdc-42* (H).

**Figure S3:**
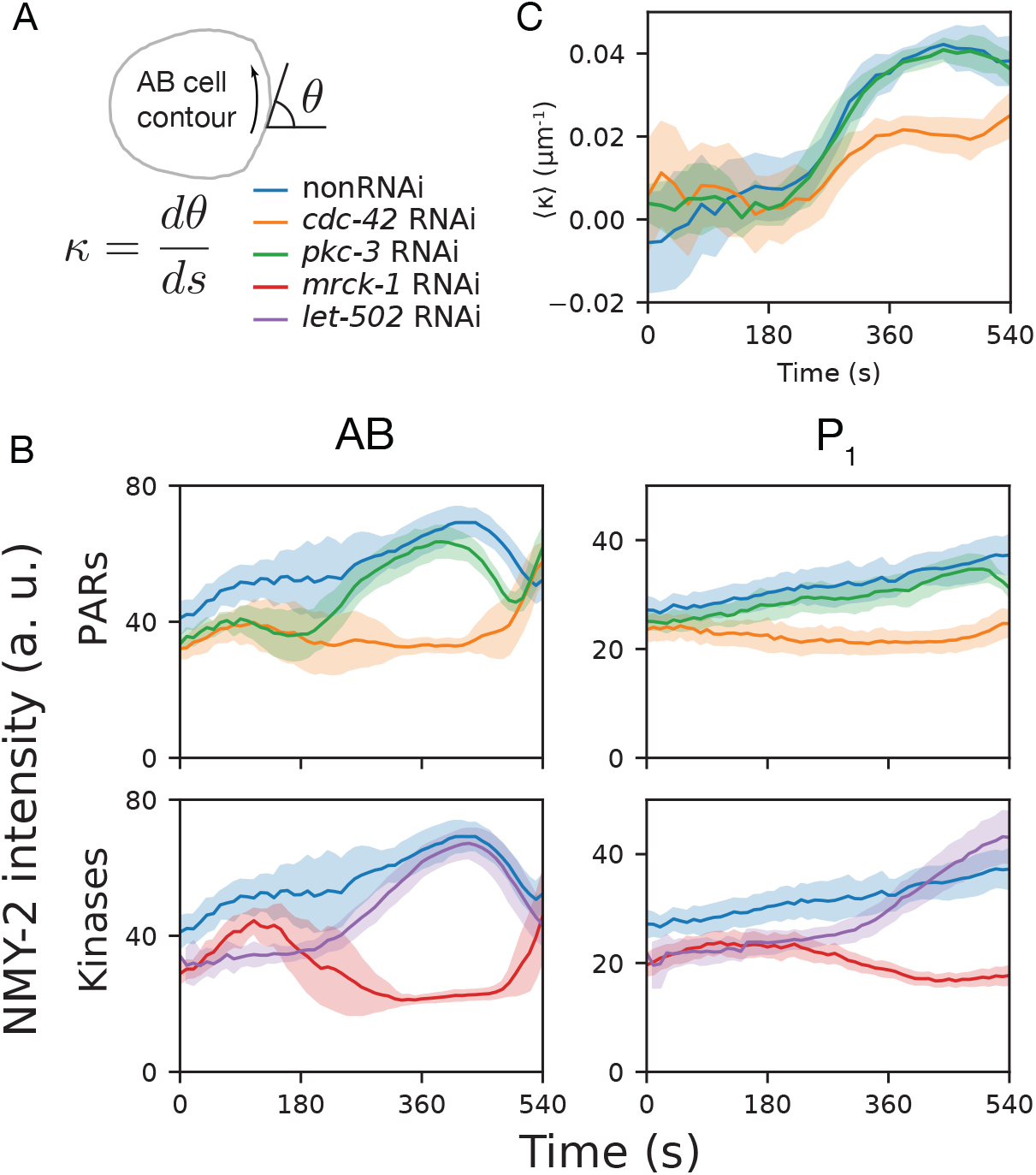
Cortical recruitment of NMY-2 is regulated by CDC-42 through MRCK-1. (A) Schematic of In-plane curvature of AB cell contour. The coordinate is counter-clockwise such that the rounded shapes give rise to positive values. (B) Cortical NMY-2 density in AB and P_1_ cells. Time T=00:00 corresponds to completion of P_0_ cytokinesis in all conditions except *let-502* RNAi embryos. Because *let-502* depletion severely slowed cytokinetic constriction in the P_0_ cell, precise alignment of the two-cell stage cell cycle was not possible using cytokinesis completion. Instead, embryos were aligned using the onset of whole-body rotation of the AB cell prior to cytokinesis. The corresponding onset time was set to T = 08:30, determined from nonRNAi embryos. (C) Time courses of in-plane curvature, spatially averaged over AB-P_1_ cell contact region, ⟨*κ*⟩.

**Figure S4:**
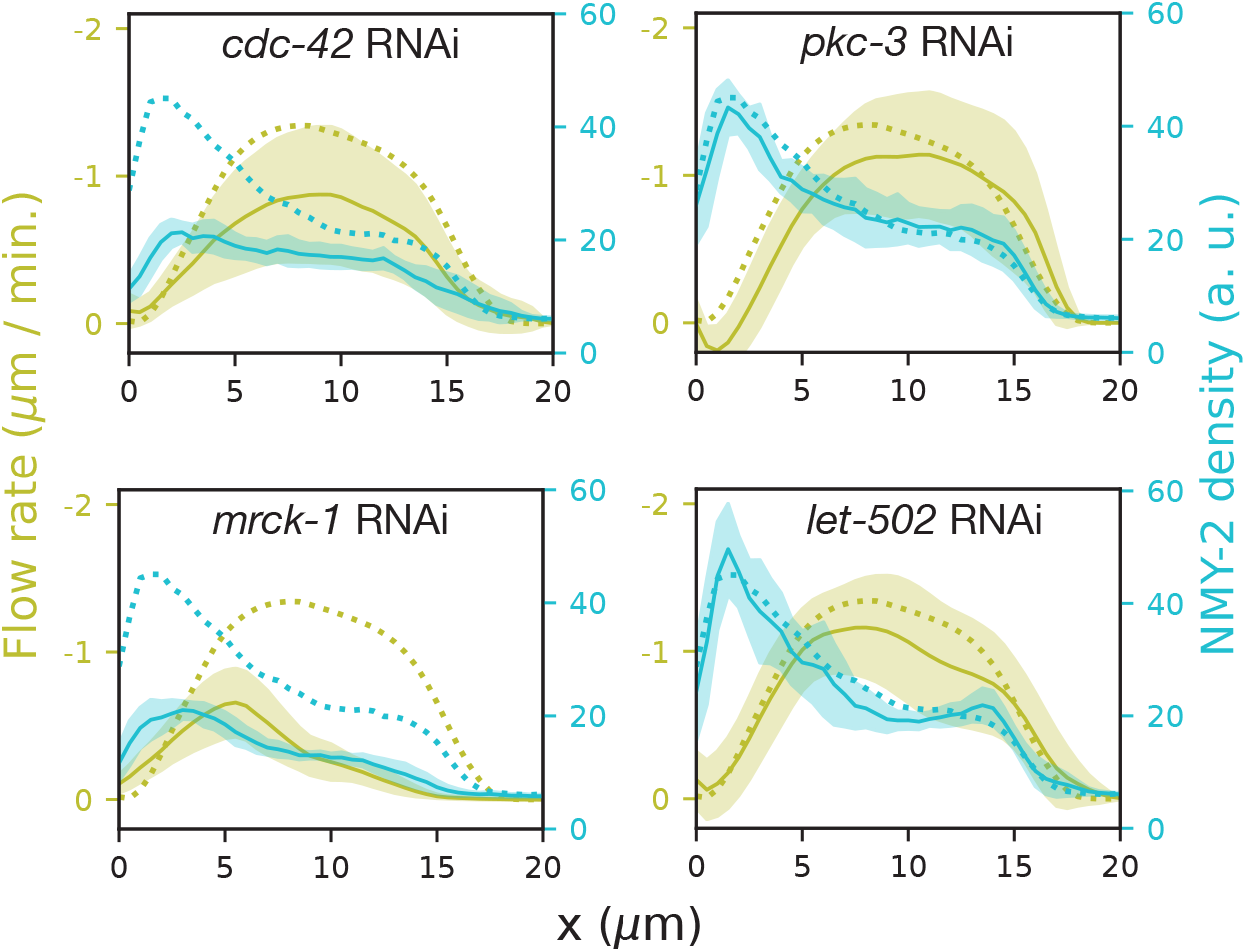
Spatial profiles of NMY-2::GFP intensity and cortical flow rate under RNAi perturbations targeting *cdc-42* (n = 10), *mrck-1* (n = 10), *pkc-3* (n = 10), and *let-502* (n = 12)

**Figure S5:**
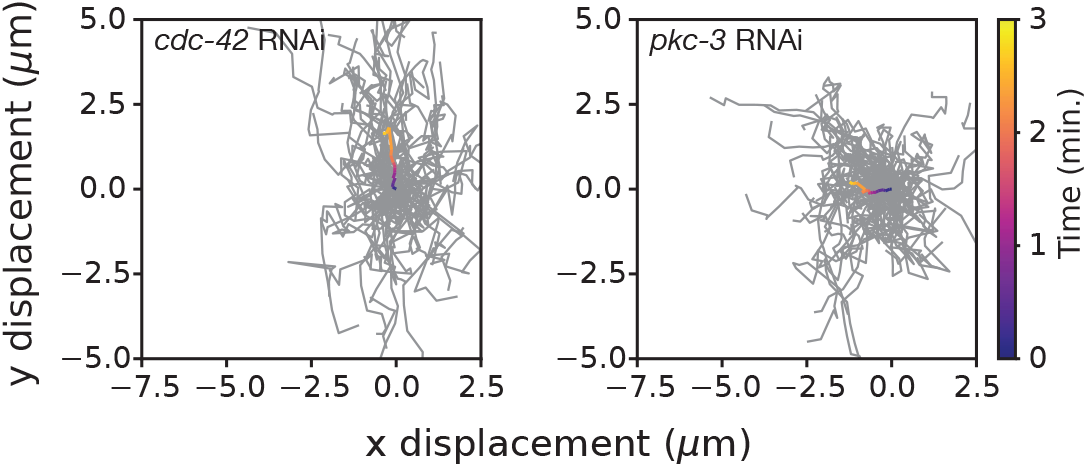
Trajectories of individual PAR-3 clusters during the polarization period. Grey lines show trajectories of individual clusters, and temporally color-coded lines indicate ensemble-averaged trajectories over time. Data are shown for *cdc-42* RNAi (upper, n = 64 clusters from 11 embryos) and *pkc-3* RNAi (lower, n = 73 clusters from 17 embryos).

